# Screening for links between behaviour and acute hyperthermia and hypoxia resistance in rainbow trout using isogenic lines

**DOI:** 10.1101/2023.10.19.563047

**Authors:** H. Lagarde, D. Lallias, F. Phocas, L. Goardon, M. Bideau, F. Guyvarc’h, L. Labbé, M. Dupont-Nivet, X. Cousin

## Abstract

In the context of adaptation to climate change, acute hyperthermia and hypoxia resistance are traits of growing interest in aquaculture. The feasibility of genetic improvement of these resistance traits through selection has been demonstrated in rainbow trout (*Oncorhynchus mykiss*). The present paper aims to test whether behaviour may be associated with acute hyperthermia and hypoxia resistance to better characterize these resistance phenotypes.

For this, six rainbow trout isogenic lines were phenotyped for behaviour variables and for acute hyperthermia and hypoxia resistance, using different individuals for each phenotype. The behaviour variables of the fish were phenotyped using an individual test in a new environment. The experimental design used 150 fish phenotyped per isogenic line for each resistance trait and 18 fish per isogenic line for behavioural traits, distributed in triplicates. Relations between acute hyperthermia and hypoxia resistance phenotypes and behaviour phenotypes were tested at the level of isogenic lines.

Significant differences in behaviour between isogenic lines were found, with some behaviour variables being highly associated with hypoxia resistance and moderately associated with acute hyperthermia resistance. Travelling distance, frequency of change between a risky and a safe zone of the tank and the percentage of time in movement in the behaviour test were strongly positively associated with acute hypoxia resistance. Travelling distance and frequency of change between a risky and a safe zone of the tank in the behaviour test were slightly negatively associated with acute hyperthermia resistance. This previously unstudied link between behaviour and resistance phenotypes also suggests that some behaviour variables could be used as predictors for acute hyperthermia and hypoxia resistance in fish. This result could lead to more ethical acute hyperthermia and hypoxia resistance phenotyping protocols, as the current protocols in use are classified as severe by French ethics committees.

## Introduction

Resistance to acute hyperthermia and hypoxia are phenotypes of interest in fish for numerous fields of research, such as evolutionary biology (Mandic, Todgham & Richards, 2009; Comte & Olden, 2017), stress-coping physiology (Rytkönen et al., 2007; Saravia et al., 2021), forecasts for climate change effects (Messmer et al., 2017; Potts, Mandrak & Chapman, 2021), and genetics for breeding robust animals (Baer & Travis, 2000; Perry et al., 2005; Debes et al., 2021; Lagarde et al., 2023a; Prchal et al., 2023). Previously, genetic component for acute hyperthermia and hypoxia resistance has been demonstrated in rainbow trout (Perry et al., 2005; Lagarde et al., 2023a,b; Prchal et al., 2023). In particular, (Lagarde et al., 2023b) showed that some genotypes were more resistant to acute hyperthermia or hypoxia using rainbow trout isogenic lines. The predominant methods in use for measuring these resistance phenotypes consist in increasing the temperature of the water for hyperthermia phenotyping or decreasing the O_2_ saturation of the water for hypoxia phenotyping until the fish lose equilibrium. The resistance phenotypes include the resistance time and the temperature/O_2_ saturation at the equilibrium loss (Beitinger, Bennett & Mccauley, 2000; Roze et al., 2013).

There is a strong interest in identifying reasonably intrusive phenotypes, that could be used as predictors of resistance to acute hyperthermia and hypoxia, to replace the current classical phenotyping methods. Indeed, although the classical phenotyping methods of acute hyperthermia and hypoxia resistance have the advantage of directly measuring the phenotypes of interest, they have drawbacks. Firstly, acute hyperthermia and hypoxia challenges push fish to their limits, with the loss of equilibrium as an endpoint generating animal suffering. Within the 3R framework (Replacement, Reduction, Refinement), active research must be conducted to reduce animal suffering in scientific experiments (Sloman et al., 2019). Secondly, they affect the physical integrity of the fish, limiting the re-use of animals for other experiments or breeding for genetic selection. For example, mortality the week following an acute hyperthermia challenge was between 5% and 55% in rainbow trout depending on the strain (Strowbridge et al., 2021). During the course of previous studies, aiming at the evaluation of the resistance to acute hyperthermia and hypoxia of rainbow trout (Dupont-Nivet et al., 2014; Lagarde et al., 2023b), we observed expression of different ranges of behaviours between lines suggesting there may be an underlying genetic component for these behaviours. Interestingly, different behaviours were previously shown to be associated with traits possibly linked with acute hyperthermia and hypoxia resistance in various fish species: proactivity was shown to be associated with thermal preference in Nile tilapia (Cerqueira et al., 2016), reactivity was linked to faster growth in brown trout *Salmo trutta* (Adriaenssens & Johnsson, 2011), and boldness in a refuge emergence test was related with high aerobic capacity in bluegill sunfish *Lepomis macrochirus* (Binder et al., 2016). In rainbow trout, resistance to acute hyperthermia was associated with lower routine activity (Campos, Val & Almeida-Val, 2018), while divergent behaviours were observed in response to hypoxia challenge, with some fish showing strong activity levels while others remained quiet, the later behaviour being associated with a higher survival rate (Van Raaij et al., 1996a). However, the research presented in these articles was not specifically designed to link behaviour to resistance to acute hyperthermia or hypoxia. In the present article, we investigate whether resistance to acute hyperthermia and hypoxia may be linked to behaviour, which was not previously done, to our knowledge. We also discuss the feasibility of using behaviour variables as a novel way of phenotyping these resistances.

Our methodology used the rainbow trout isogenic lines, which have been previously shown to have different resistance to either hyperthermia or hypoxia (Lagarde et al., 2023b). The behaviour of fish was phenotyped in an individual behaviour test. The measured behaviour phenotypes were describing activity (maximum acceleration, maximum velocity, distance travelled, movement) and risk-taking (emergence, time spent in a risky zone). The resistance and behaviour phenotypes, measured on different individuals, were related at the level of lines, thanks to the complete genetic homogeneity among fish within an isogenic line. Based on our previous knowledge of coincidental differences in both behavioural traits and hyperthermia and hypoxia resistance in different rainbow trout lines, we hypothesised that there may be correlations between both. If this were the case, this would provide an operational and less intrusive test which could be useful in selection programs.

## 1 Material and methods

In the present study, three phenotypes were collected on six isogenic lines: acute hyperthermia resistance, acute hypoxia resistance and individual behaviour. Rearing of fish methods and phenotyping protocols are presented below.

### 1.1 Experimental fish

Experimental fish were produced at the PEIMA (Pisciculture Expérimentale INRAE des Monts d’Arrée) experimental fish farm (doi: 10.15454/1.5572329612068406E12, Sizun, France) with broodstock issued from the INRAE (French national research institute for agriculture, food and environment) homozygous isogenic rainbow trout lines (Quillet et al., 2007). Homozygous isogenic lines are a powerful genetic resource, as fish are genetically identical within a line. Therefore, fish from a given line can be considered as replicates of a unique genotype.

The fish used in the present study are from the same rearing batch as those described in (Lagarde et al., 2023b). Figure 1 summarises the rearing process of the fish and the experiments conducted. Briefly, six heterozygous isogenic lines were produced by fertilising eggs from same-day naturally spawning females of the homozygous isogenic line B57 with the sperm of six neomales (sex-reversed XX males) issued from the homozygous isogenic lines A02, A22, A32, B45, N38 and R23. At this stage, the only source of variation between heterozygous lines was expected to be the paternal genetic effect. Fertilised eggs were incubated in one hatching tray per line, supplied with spring water of little thermal amplitude over the year (11.5°C - 12.4°C). At 21 dpf (days post-fertilisation), hatching trays were each randomly divided into three hatching trays with no mixing between isogenic lines (>500 eyed eggs per hatching tray, 3x6 hatching trays). At 43 dpf, each hatching tray was transferred into a 0.25 m^3^ indoor tank with no mixing between hatching trays (∼500 larvae per tank, 3x6 tanks) supplied with the same spring water. At 145 dpf, the number of fries was reduced to 230 in each of the 18 tanks by randomly removing exceeding fries. At 150 dpf, the number of individuals was randomly reduced to 160 in each of the eighteen tanks, decreasing the density from 7.0 to 4.5 kg/m^3^. The same day, among the approximately 160 kept individuals per tank, 100 were individually anaesthetised, PIT-tagged (Biolog-id, pit-tag dimensions: 12*9*2 mm, weight: 0.1g) in the dorsal muscle just behind the head, and put back into their tanks.

**Fig. 1.**
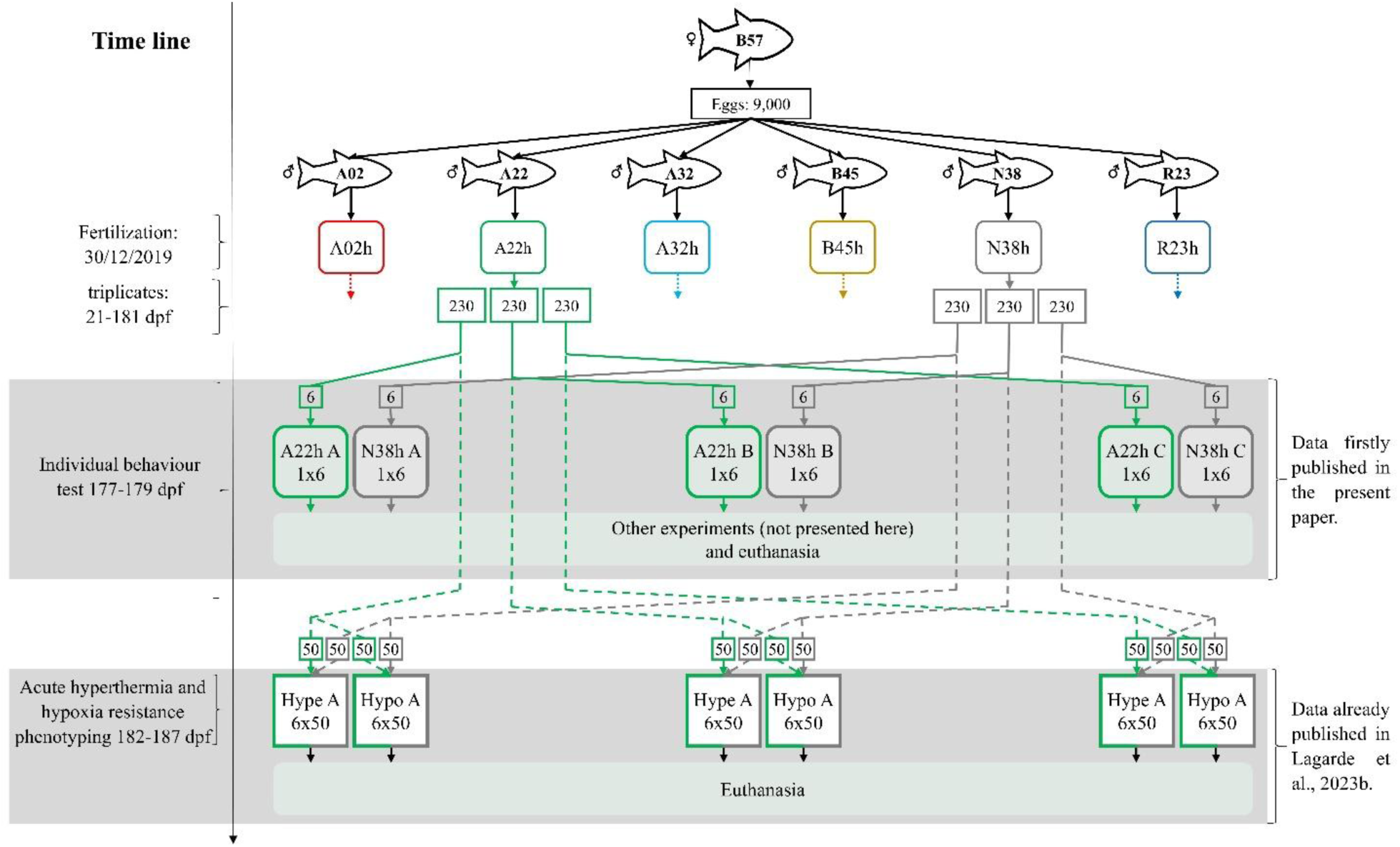
Rearing history of fish and experimental design. Each isogenic line is represented by a different colour. The entire rearing history and experimental use were only represented for lines A22h and N38h. The four other lines were not represented to not overload the figure, but they had a similar life history and were phenotyped following the same protocols. Rounded rectangles represent rearing tanks. Replicate rearing tanks also corresponding to replicate behaviour and resistance phenotyping are named A, B and C. The theoretical number of fish is indicated inside each rectangle. The number of fish and lines used in each experiment is indicated in each experiment rectangular, with the first number indicating the number of isogenic lines and the second the number of fish per isogenic line. The lines represent fish movements for phenotyping, solid for fish used in behaviour test and dotted for fish used in acute hyperthermia and hypoxia challenges. "Hype" stands for acute hyperthermia phenotyping, and "Hypo" for acute hypoxia phenotyping. Figure adapted from Lagarde et al. (2023b).

At 175 dpf, PIT-tagged fish were removed from rearing tanks to form the three replicates for phenotyping the resistance of isogenic lines to acute hyperthermia and the three replicates for phenotyping the resistance of isogenic lines to acute hypoxia. The resistance to acute hyperthermia and hypoxia were phenotyped between 182 and 187 dpf as already presented in Lagarde et al. (2023b).

Among the 60 untagged fish kept in the rearing tanks, six were randomly collected to evaluate individual behaviour between 177 and 179 dpf according to experimental setups presented below.

During the rearing process, the maximum density was 8.7 kg/m^3^, and fish were fed to satiation with a commercial diet from Le Gouessant’s company.

### 1.2 Conduct of acute hyperthermia and hypoxia resistance tests

The methods and devices used to determine the mean acute hyperthermia and hypoxia resistance of isogenic lines were previously presented in (Lagarde et al., 2023b) and will be briefly summarized here.

Acute hyperthermia and hypoxia challenges were performed between 182 dpf and 187 dpf. Fish mean body weight was 15.5 ± 3.5 g. Isogenic lines were phenotyped in triplicate for acute hyperthermia and hypoxia resistance (Figure 1). Each replicate (named A, B and C in Figure 1) consisted of 300 fish, composed of 50 fish from each of the 6 isogenic lines. The evening prior to each challenge, fish of the challenged replicate were moved to the indoor challenge tank (0.12 m3) supplied with the same water source as the one used in the respective rearing tanks and left alone for the night for acclimation (Lagarde et al 2023b).

For the acute hyperthermia challenge, the temperature in the challenge tank was first increased at a rate of 3.6°C/hour (12.4°C to 23.2°C in three hours). The temperature increase rate was then reduced to 0.6°C/hour and maintained at this rate until all fish lost equilibrium. For comparison, in rainbow trout, the optimum temperature is 16-17°C for growth and 10-13°C for reproduction (MacIntyre et al., 2008). O_2_ level was controlled by bubbling pure O_2_ (Lagarde et al., 2023b). The temperature increase rate was manually controlled by adding water from a heated buffer tank.

For the acute hypoxia challenge, O_2_ saturation in the challenge tank was decreased from nearly 90% to 15% in 4.5 hours, by bubbling dinitrogen gas into a saturation column (Lagarde et al 2023b). O_2_ saturation was then stabilised around 15% for the rest of the challenge. For comparison, in rainbow trout, at 13°C for 28 days, a drop of O_2_ saturation from 90% to 40% decreased the survival and growth by 7% and 10%, respectively (Jiang et al., 2021). In acute hypoxia challenge, the temperature remained in a range suitable for rainbow trout (10-14°C). Temperature and O_2_ concentration were recorded every 5 min during acute hyperthermia and hypoxia challenges using electronic probes (HQ40d, Hach Company, Loveland, CO, USA) (Lagarde et al 2023b).

As temperature increased in acute hyperthermia challenge or O_2_ saturation decreased in acute hypoxia challenge, fish gradually lost equilibrium. Immediately after a fish lost equilibrium (defined as an inability to maintain upright swimming for 10 seconds), it was caught, its isogenic line was identified by reading the PIT tag, and the time was recorded. The fish was then anaesthetised (Tricaïne MS222, 50 mg/L), weighed, and euthanised by an overdose of anaesthetic (Tricaïne MS222, 150 mg/L).

Acute hyperthermia resistance was quantified as the cumulative thermal exposure (CTE) in degree-minutes (deg-min), and acute hypoxia resistance was quantified as the cumulative hypoxia exposure (CHE) in saturation minutes (sat-min), as described in Lagarde et al. (2023b). CTE is the sum of the differences between the temperature in the challenge tank and the temperature at the start of the challenge, calculated at each minute from the start of the challenge to the time of loss of equilibrium. CHE is the sum of the differences between a reference O_2_ saturation of 90%, the O_2_ saturation before the start of the challenge, and the O_2_ saturation measured in the challenge tank, calculated at each minute from the start of the challenge to the time of loss of equilibrium. A high CTE or CHE indicates a high resistance to acute hyperthermia or hypoxia, respectively.

As explained in Lagarde et al. (2023b), the effect of isogenic line on acute hyperthermia and hypoxia resistance was analysed by fitting a model with *line* as a fixed effect, fish *body weight* as a covariate, their interaction, and replicate as a random effect. *line*, *body weight* and their interaction were all significant in hyperthermia and hypoxia challenges (Lagarde et al. 2023b). Function lme in package nlme 3.1–153 (Pinheiro et al., 2021) was used to fit linear mixed models. Because the effect of *body weight* on resistance was different between lines (P<0.05), the predicted lsmeans of *lines* on acute hyperthermia and hypoxia resistance were estimated at 15.5 g, the mean body weight, using lsmeans 2.30–0 R package (Lagarde et al. 2023b).

### 1.3 General conduct of behavioural test

The purpose of the individual behaviour test was to evaluate behaviour in a novel environment. Behavioural test was carried out in a quiet room with no traffic during the experiments. Figure 2 presents the diagram of the experimental devices.

**Fig. 2.**
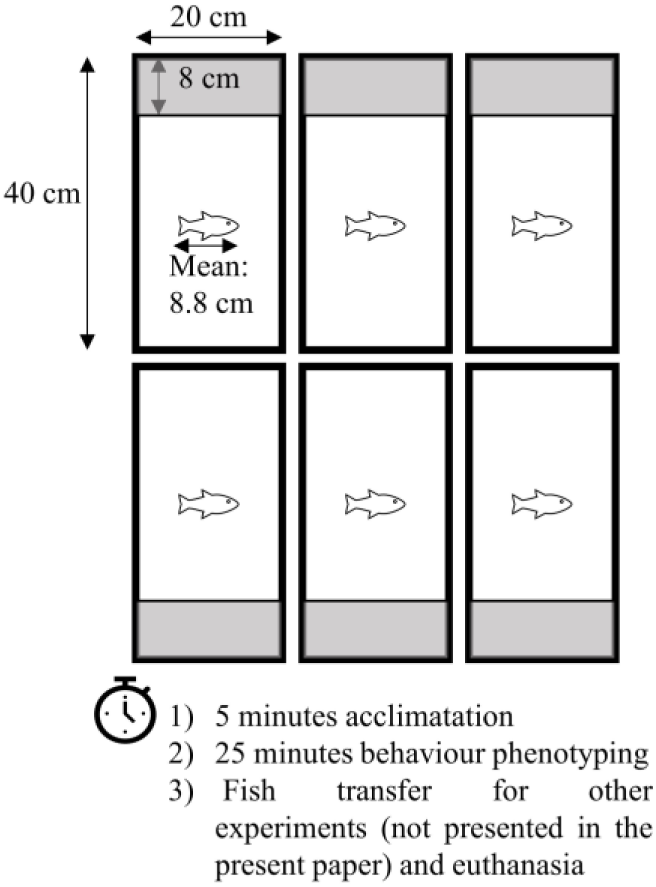
Diagram of behaviour test. Grey bands represent the safe zone, covered with a lid opaque to visible light. The six fish monitored in parallel were from the same isogenic line.

Behaviour was simultaneously evaluated on six isolated individuals from the same isogenic line for 25 minutes. Devices and duration were derived from previously published protocols used on zebrafish and seabass (Vignet et al., 2015; Alfonso et al., 2020; Alfonso et al., 2020; Alfonso, Gesto & Sadoul, 2021). The same water source with similar temperature was used in home tanks and during the behaviour test. Behaviour tests were conducted in triplicates for each of the six lines, totalling 18 tests. Each test lasted approximately 30 minutes: 5 minutes of acclimation and 25 minutes of behavioural test. Fish were finally anaesthetized (Tricaïne MS222, 50 mg/L), weighed using digital scales, and euthanized by an overdose of anaesthetic (Tricaïne MS222, 150 mg/L). Behaviour tests were spread over three days at a rate of 6 sessions per day. They started at six fixed different hours in the day: 9h00, 10h00, 11h00, 13h00, 14h00, 15h00. We arranged sessions for the three replicates of each isogenic line during morning, midday and afternoon slots so that there was no confounding between line and time slot.

Behaviour test started by catching six fish from the same isogenic line in one of the rearing tanks according to the pre-determined session order. Fish were then immediately placed randomly in six individual rectangular aquariums (20×40 cm, Europrix) filled with 12 L of water corresponding to 15 cm depth of water (Figure 2). The six aquariums were placed in a homemade rectangular tent lit by spots on the outside top of the tent to ensure homogeneous semi-darkness conditions. Aquariums were isolated from neighbouring aquariums by opaque white walls and backlit by an infrared device (IR floor 1×1 m, Noldus, The Netherlands) as infrared radiation is invisible for trout (Rader et al., 2007).

Aquariums were divided into two zones: on one side, a zone of size 20 x 8 cm covered by a plate opaque to visible light but transparent to infrared light called the safe zone, and on the other side, an uncovered zone of size 20 x 32 cm called the risky zone (Figure 2 and actual video capture as Fig. S1). Fish were first placed in the safe zone, separated from the risky zone by a door opaque and completely closing the space between the safe and risky zones. Fish were left for 5 minutes in the safe zone, and the door was then lifted using an electric engine remotely controlled allowing fish to freely move between the safe and risky zones. The two engines, each lifting the three doors of the three contiguous tanks, were located on a structure independent from the tanks to prevent vibration and were activated simultaneously to ensure synchrony of the opening. Once the door opened, fish activity was video recorded for 25 minutes with a USB camera (DMK 31AU03, The Imaging Source) equipped with a 2.7–13.5-mm lens (Fujinon) and an infrared filter (The Imaging Source), using ICapture (The Imaging Source). After 25 minutes, the fish were immediately caught and placed in a bucket filled with same water. At the end of each test, water of aquariums was entirely renewed before the start of the following test.

### 1.4 Videos’ behaviour variables extraction

Videos were analysed with EthoVision XT15.0 software. Fish positions were tracked using a dynamic subtraction method. The accuracy of the tracked positions was checked by viewing the superposition of the tracked and actual positions of fish in all videos and manually editing if required. The Lowess smoothing method was applied to eliminate background noise.

Eight behaviour variables were measured: the maximum acceleration (ACC_MAX, cm s^-2^), the distance travelled (DIST_TRAV, cm), the latency to first exit from the safe zone (EMERGENCE, s), the zone change frequency (FRQ_CHAN, # min^-1^), the absolute meander (MEANDER, deg cm^-1^), the moving duration (MOV%, % of the time), the time spent in the risky zone (RISK%, % of the time) and the maximum velocity (VEL_MAX, cm s^-1^) (Table 1). To consider the effect of time on behaviours, behaviour variables were extracted over several time bins, every 5 minutes. This was done to evaluate the potential effects of acclimation of the fish along the duration of the behaviour test.

**Table 1.** Definition of behaviour variables.

| Abbreviation | Full name | Unit | Details |
| --- | --- | --- | --- |
| ACC_MAX | Maximum acceleration | cm.s<sup>-2</sup> | The maximum acceleration. Acceleration is the difference in velocity divided by the time difference between the current sample and the previous one. |
| DIST_TRAV | Distance moved | m | The cumulated distance travelled by the body centre point of the subject from the previous sample to the current one. |
| EMERGENCE | Latency to exit from the safe zone | s | The time until fish first exit from the safe zone |
| FRQ_CHAN | Zone change frequency | #.min<sup>-1</sup> | The number of time per minute the subject cross the border between the risky and the safe zone. |
| MEANDER | Meander | deg.cm<sup>-1</sup> | Meander is the change in direction of movement of a subject relative to the distance moved by that subject |
| MOV% | Percentage of the time moving | % | The fraction of time the subject was in moving state, *i.e.* when the subject velocity was above 1.5 cm/s. |
| RISK% | Percentage of the time in the risk zone | % | The fraction of time the subject remains in the risky zone. |
| VEL_MAX | Maximum velocity | cm.s<sup>-1</sup> | The maximum velocity. Velocity is the distance moved by the body centre point of the subject per unit of time. The velocity was averaged over 2 frames. |

Body weight was found to have a significant effect on acute hyperthermia and hypoxia resistances in our experiment, and resistance of isogenic lines was corrected for this covariate (Lagarde et al. 2023b). It was therefore necessary to take the fish size effect into account when correcting behaviour phenotypes. Body weight of fish used in behaviour test was measured individually but after regrouping the fish in a bucket at the end of the behaviour test. It was therefore not possible to assign each body weight to each individual fish. Fortunately, in rainbow trout, fork length is an accurate predictor of body weight, with a phenotypic correlation of 0.9 (Haffray et al., 2013; Lagarde et al., 2023a). Fork length of each fish was measured *a posteriori* by image analysis of the video recorded using ImageJ software (https://imagej.net).

Measurement of the length of each fish was repeated three times per fish, on a similar number of different screenshots with fish in a straight position. Scaling was carried out using the known length of the tank. Coefficient of variation between the 3 different measurements of an individual fish was under 3 %, ensuring a good reliability on length measurement.

### 1.5 Behaviour data statistical analysis

Analyses were performed in R version 4.0.3 (R Development Core Team, 2019).

Behavioural variables were measured individually (6 isogenic lines x 3 replicas x 6 individuals, 108 data points per behaviour variable and time bin). The experimental unit was the individual fish.

Mean fork length differences among isogenic lines were analysed by analysis of variance (ANOVA), using stats package. ANOVA assumptions of normality and homoscedasticity were verified by visual inspection of residual-fit plots.

To test for an effect of the isogenic line through time bins on behaviour variables, we fitted linear mixed effects models with *fork length* as a covariate, *time* (time bins) and *line* (isogenic line) as categorical fixed effects, *replica* as a random effect, and the interactions between *time*, *line* and *fork length*. Time was set up as a categorical variable because a visual inspection of response variables plotted against time did not reveal any linear relationship. For EMERGENCE, only *line* and *fork length* effects were tested, as emergence can only happen once in the whole test. Function lme in package nlme 3.1–153 (Pinheiro, J. et al., 2021) was used to fit linear mixed models. Model selection (with vs. without *time* and/or *line* and/or *fork length* and/or the different interactions between these variables) was done using the stepAIC function in the MASS package (Venables & Ripley, 2002). In a second step, weighted terms allowing variances to differ among isogenic lines were considered, choosing the model with the lowest AIC (with vs. without weighted terms) (Zuur et al., 2009). To account for temporal autocorrelation, using package nlme 3.1–153 (Pinheiro, J. et al., 2021), autoregressive AR1/AR2 or autoregressive moving average ARMA(1,1)/ARMA(2,1) covariance structures were considered, choosing the model with the lowest AIC (AR1/AR2/ARMA(1,1)/ARMA(2,1)) structures (Zuur et al., 2009). Assumptions of normality, homoscedasticity and independence of final linear mixed models were verified by visual inspection of residual-fit plots. The significance of model effects was tested using F-tests with package lmerTest 3.1–3.

The predicted least-squares means (lsmeans) of *line* and *time* on each response variable were estimated using lsmeans 2.30–0 R package (https://cran.r-project.org/package=lsmeans). For behaviour variables for which the *fork length* effect was significant, lsmeans of *time* and *line* (when significant) were estimated considering a fork length of 8.8 cm, corresponding to the mean fork length of fish used in the behaviour experiment. Differences among *line* and *time* lsmeans for behaviour variables were compared by Tukey’s test with Type 1 error risk set at 0.05 using lsmeans 2.30–0 R package (Lenth & Love, 2015). For behaviour variables for which the best model included the *line* × *time* interaction, *line* lsmeans were calculated at each *time* bin and vice-versa.

### 1.6 Screening for relationships between behaviour and resistance variables

The lsmeans of the behavioural variables and the resistance to acute hyperthermia and hypoxia stress were calculated for the same six isogenic lines using the steps mentioned in the previous sections. Pearson correlation coefficients between lsmeans of lines for behavioural variables and for resistance to stress were calculated using stats 3.6.2 R package. Behaviour variables for which lsmeans of isogenic lines were highly correlated with lsmeans of isogenic lines on resistance phenotype were considered as potentially genetically linked with resistance phenotypes. The significance of correlations with Type 1 error risk set at 0.05 was estimated using the function cor.test of the package stats.

## 2 Results

### 2.1 Resistance phenotypes

There were significant differences in acute hyperthermia and hypoxia resistance between isogenic lines (Table 2). This result is presented in more details and discussed in (Lagarde et al., 2023b).

**Table 2.**
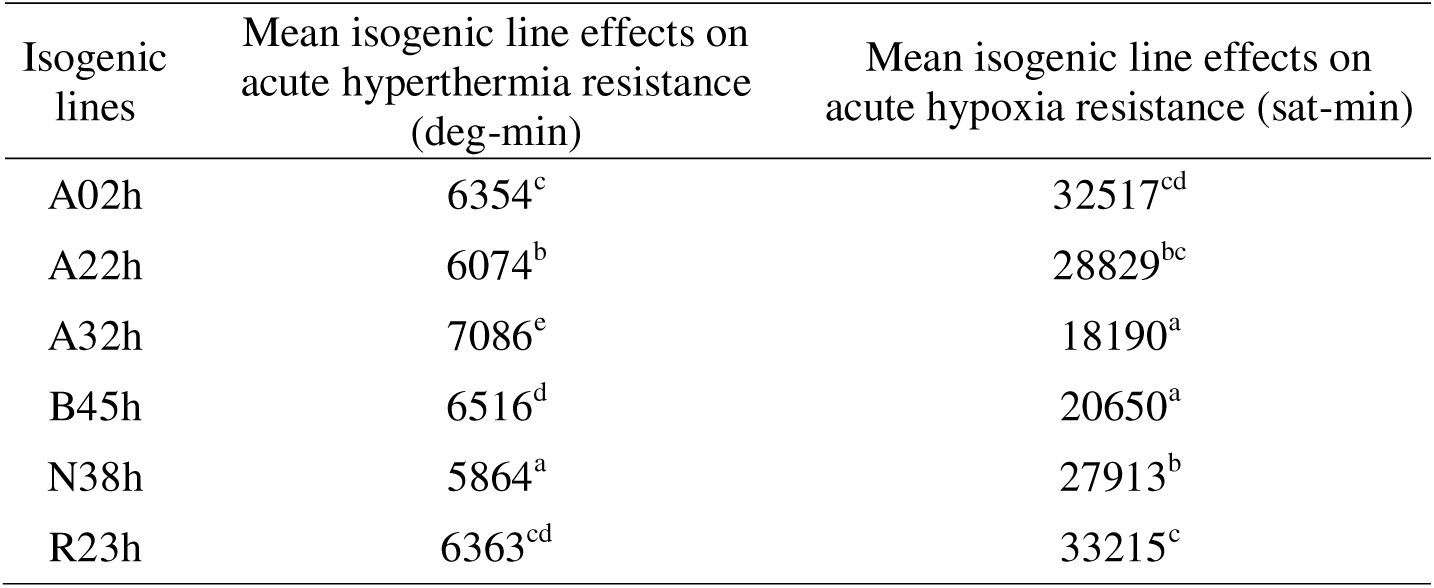
Isogenic lines means (lsmeans) for acute hyperthermia resistance and acute hypoxia resistance. Means with same letters within columns are not significantly different.

### 2.2 Fork length and body weight in resistance and behaviour experiments

In behaviour test, the mean fish body length and weight were 8.8 ± 1.0 cm and 11.6 ± 3.2 g, respectively. There were significant differences in fork length between lines (ANOVA, N = 108, df=5; F value = 9.6; P < 1.6e-07) and body weight (ANOVA, N = 108, df=5; F value = 17.3; P < 2.8e-14). The smallest/lightest line was A32h (8.2 ± 0.6 cm, 9.6 ± 2.6 g), and the tallest/heaviest was N38h (9.9 ± 0.5 cm, 14.9 ± 2.8 g) (Table S1). At the isogenic line level, the Pearson coefficient of correlation between mean fork length and mean body weight was 0.99 (not shown), confirming that fork length is a good proxy for body weight.

In resistance challenges, mean body weight was 15.5 ± 3.5 g. There were also significant differences in body weight between lines (Lagarde et al. 2023b). The fork length was not measured in resistance challenges. The lightest line was A32h (12.5 ± 2.6 g), and the heaviest was N38h (18.3 ± 2.8 g) (Table S1). Mean body weight was lower in the behaviour tests (11.6 ± 3.2 g) than in resistance challenges (15.5 ± 3.5 g) since fish had between one and two more weeks to grow in the resistance challenges. Rankings of lines based on body weight were similar between behaviour test and resistance challenges (Figure S2).

### 2.3 Factors influencing behaviour variables

The significance of effects (*line*, *time*, *fork length* and their interactions) on the different behaviour variables is shown in Table 3. Details about the F-statistic value and degrees of freedom of the numerator and denominator are given in Table S2. *isogenic line* effect was significant for all studied behaviour variables except EMERGENCE (Table 3). *time* effect was only significant for ACC_MAX and VEL_MAX (Table 3). *length* was never significant without interaction with *line* (Table 3). The only significant interactions were *line* × *length* interactions for ACC_MAX, MOV%, RISK% and VEL_MAX. For these variables, *isogenic line* effect was estimated at the mean body length, *i.e.* 8.8 cm.

**Table 3.** Best model for behaviour variables. *line: isogenic line; time: time bin; length: fork length, AR: AutoRegressive; ARMA: AutoRegressive Moving Average.* In F-test, “*” means the P-value is inferior to 0.05 significant threshold. according to AIC test.

| Behaviour variables | Fixed structure best model<sup>1</sup> | Weighted terms for line effect<sup>1</sup> | Temporal autocorrelation<sup>1</sup> |
| --- | --- | --- | --- |
| ACC_MAX | *line*<sup>*</sup>, *time*<sup>*</sup>, *length*<sup>*</sup>, *line*×*length*<sup>*</sup> | yes | no |
| DIST_TRAV | *line*<sup>*</sup>, *length* | yes | AR(2) |
| EMERGENCE | no significant effect | no | no |
| FRQ_CHAN | *line*<sup>*</sup>, *length* | yes | ARMA(1) |
| MEANDER | *line*<sup>*</sup> | yes | AR(1) |
| MOV% | *line*<sup>*</sup>, *length*, *line*×*length*<sup>*</sup> | yes | AR(2) |
| RISK% | *line*<sup>*</sup>, *length*<sup>*</sup>, *line*×*length*<sup>*</sup> | no | ARMA(2,1) |
| VEL_MAX | *line*<sup>*</sup>, *time*<sup>*</sup>, *length*, *line*×*length*<sup>*</sup> | yes | AR(2) |

EMERGENCE was the only behaviour variable for which neither *time*, *line* nor *fork length* effects were significant (Table 3). This variable was therefore not investigated further. Regression coefficients of fork length on behaviour variables are presented in Table S3. Effect of size on maximum acceleration was positive for A32h, N38h and R23h and negative for A02h, A22h and B45h. For all isogenic lines except A32h, taller fish have smaller percentage of the time moving (MOV%) and maximum velocity (VEL_MAX) compared to smaller fish (Table S3). For A32h line, the relationship is opposite with smaller fish having smaller percentage of the time moving (MOV%) and maximum velocity (VEL_MAX) compared to taller fish. Eventually, fork *length* has a negative effect on RISK% for lines A22h, N38h and R23h and a positive effect for lines A02h, A32h and B45h (Table S3).

The *time* effect influenced ACC_MAX and VEL_MAX (Table 3). *time* effect and *isogenic line* were independent whatever the behaviour variable (Table 3). The maximum acceleration and the maximum velocity significantly decreased between the first and the last time bin, reflecting change in behaviour over time (Table S4).

The *isogenic line* effect influenced ACC_MAX, DIST_TRAV, FRQ_CHAN, MEANDER, MOV%, RISK% and VEL_MAX (Table 3). A02h tend to have the highest DIST_TRAV, FRQ_CHANGE, MOV% and VEL_MAX compared to other lines (Table 4) and A32h and B45h tend to have the lowest DIST_TRAV, FRQ_CHANGE, MOV% and VEL_MAX (Table 4).

**Table 4.** Isogenic lines means (lsmeans) for behaviour variables for which *line* effect was significant. Means with same letters within columns are not significantly different. For ACC_MAX, MOV%, RISK% and VEL_MAX, the *line* × *length* interaction was significant (see Table 3); therefore, isogenic line effect was estimated at the mean body length of 8.8 cm.

| Line | ACC_MAX<br>(cm.s<sup>-2</sup>) | DIST_TRAV<br>(m) | FRQ_CHANGE<br>(#.min<sup>-1</sup>) | Meander<br>(deg.cm<sup>-1</sup>) | MOV%<br>(%) | RISK%<br>(%) | VEL_MAX<br>(cm.s<sup>-1</sup>) |
| --- | --- | --- | --- | --- | --- | --- | --- |
| A02h | 38.9<sup>ab</sup> | 3.4<sup>c</sup> | 0.9<sup>c</sup> | 13.1<sup>a</sup> | 17.3<sup>c</sup> | 71.7<sup>b</sup> | 17.6<sup>c</sup> |
| A22h | 43.0<sup>b</sup> | 2.3<sup>bc</sup> | 0.7<sup>bc</sup> | 15.3<sup>a</sup> | 7.1<sup>ab</sup> | 32.9<sup>a</sup> | 17.2<sup>bc</sup> |
| A32h | 29.5<sup>ab</sup> | 0.8<sup>a</sup> | 0.2<sup>a</sup> | 16.6<sup>ab</sup> | 3.7<sup>a</sup> | 57.9<sup>ab</sup> | 12.0<sup>abc</sup> |
| B45h | 23.6<sup>a</sup> | 1.4<sup>ab</sup> | 0.4<sup>ab</sup> | 17.2<sup>a</sup> | 6.3<sup>ab</sup> | 47.7<sup>ab</sup> | 10.0<sup>a</sup> |
| N38h | 39.4<sup>ab</sup> | 2.7<sup>bc</sup> | 0.6<sup>abc</sup> | 32.3<sup>ab</sup> | 9.8<sup>abc</sup> | 29.3<sup>a</sup> | 16.3<sup>abc</sup> |
| R23h | 28.0<sup>ab</sup> | 2.3<sup>bc</sup> | 0.6<sup>bc</sup> | 39.4<sup>b</sup> | 13.8<sup>bc</sup> | 56.6<sup>ab</sup> | 11.8<sup>ab</sup> |

### 2.4 Relationships between behaviour variables

Correlations between mean effect of isogenic lines (lsmeans) for behaviour variables are presented in Table 5. DIST_TRAV, FRQ_CHANGE and VEL_MAX were highly positively correlated together with a correlation coefficient (r) ranging from 0.77 to 0.98 (Table 5). ACC_MAX was highly correlated with VEL_MAX with a r of 0.98 and moderately with DIST_TRAV and FRQ_CHANGE with a r from 0.65 to 0.66 (Table 5). MOV% was highly positively correlated with DIST_TRAV and FRQ_CHANGE (r from 0.83 to 0.87) (Table 5). RISK% was not strongly correlated with any of the other variables (r from -0.34 to 0.48) (Table 5).

**Table 5.**
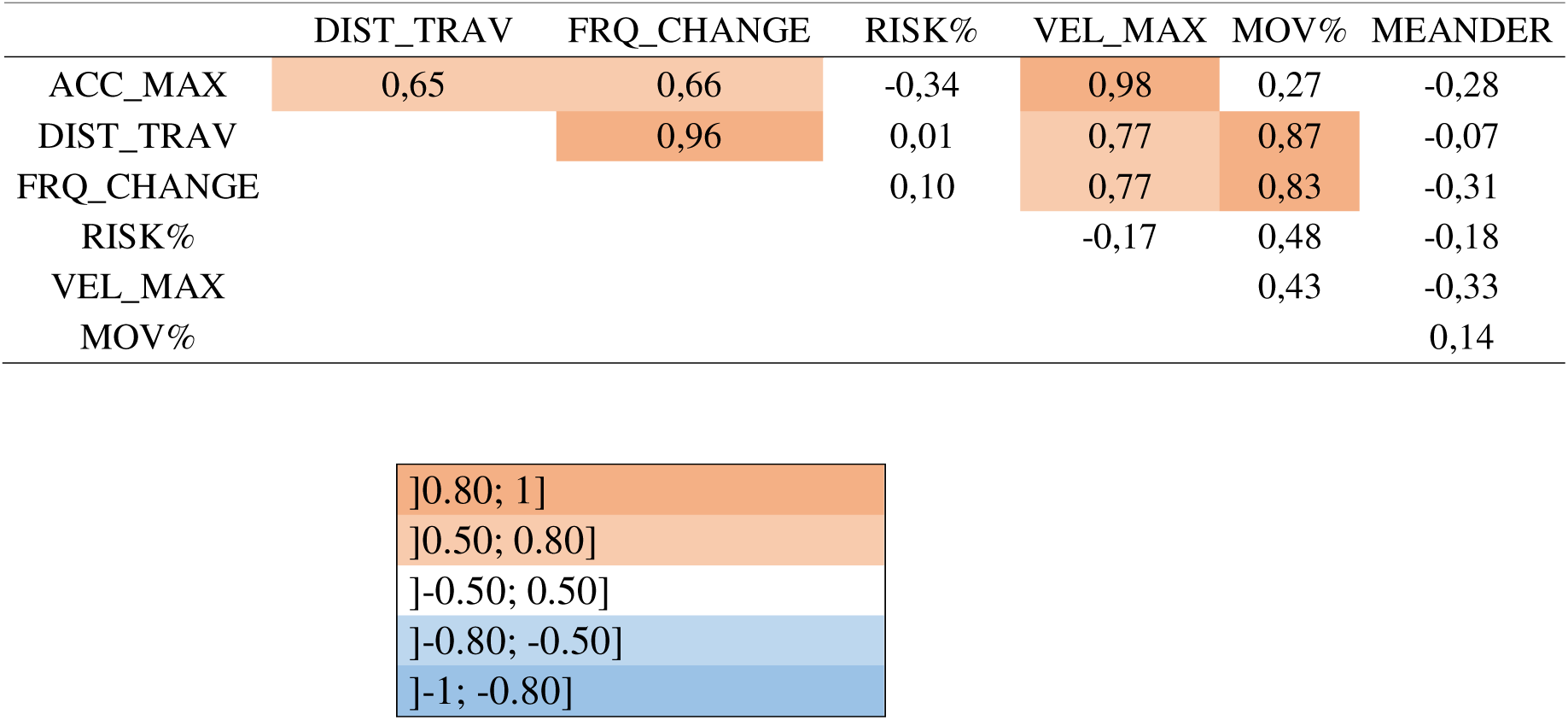
Correlations between mean effect of isogenic lines (lsmeans) for behaviour variables for which there was a significant *line* effect (see Table 3).

|  | DIST_TRAV | FRQ_CHANGE | RISK% | VEL_MAX | MOV% | MEANDER |
| --- | --- | --- | --- | --- | --- | --- |
| ACC_MAX | 0,65 | 0,66 | -0,34 | 0,98 | 0,27 | -0,28 |
| DIST_TRAV |  | 0,96 | 0,01 | 0,77 | 0,87 | -0,07 |
| FRQ_CHANGE |  |  | 0,10 | 0,77 | 0,83 | -0,31 |
| RISK% |  |  |  | -0,17 | 0,48 | -0,18 |
| VEL_MAX |  |  |  |  | 0,43 | -0,33 |
| MOV% |  |  |  |  |  | 0,14 |

|  |
| --- |
| ]0.80; 1] |
| ]0.50; 0.80] |
| ] -0.50; 0.50] |
| ] -0.80; -0.50] |
| ] -1; -0.80] |

### 2.5 Relationships between behaviour variables and resistance phenotypes

The relationships between behaviour variables and resistance to acute hyperthermia are presented on Figure 3. A32h and B45h, the two most resistant lines to acute hyperthermia, tend to have lower DIST_TRAV and FRQ_CHAN behaviour variables compared to the four other lines (Figure 3 B&C). A22h and N38h, the most sensitive lines to acute hyperthermia, tend to have smaller RISK% compared to the four other lines (Figure 3 F). However, the absolute values of the correlation coefficients between these three behaviour variables and resistance to acute hyperthermia remain moderate, between 0.61 and 0.74. This suggests a possible, but probably weak, link between these behaviour variables and resistance to acute hyperthermia. The relationships between behaviour variables and resistance to acute hypoxia are presented on Figure 4. There was a high correlation between DIST_TRAV, FRQ_CHAN and MOV% behaviour variables and acute hypoxia resistance with a r between 0.88 and 0.92 (Figure 4 B&C&E). This suggests a possible, probably strong, link between these behaviour variables and resistance to acute hypoxia. Behaviour variables whose relationship with resistance phenotypes was not mentioned are probably not linked with resistance phenotypes given the reranking observed and the low correlation.

**Fig. 3.**
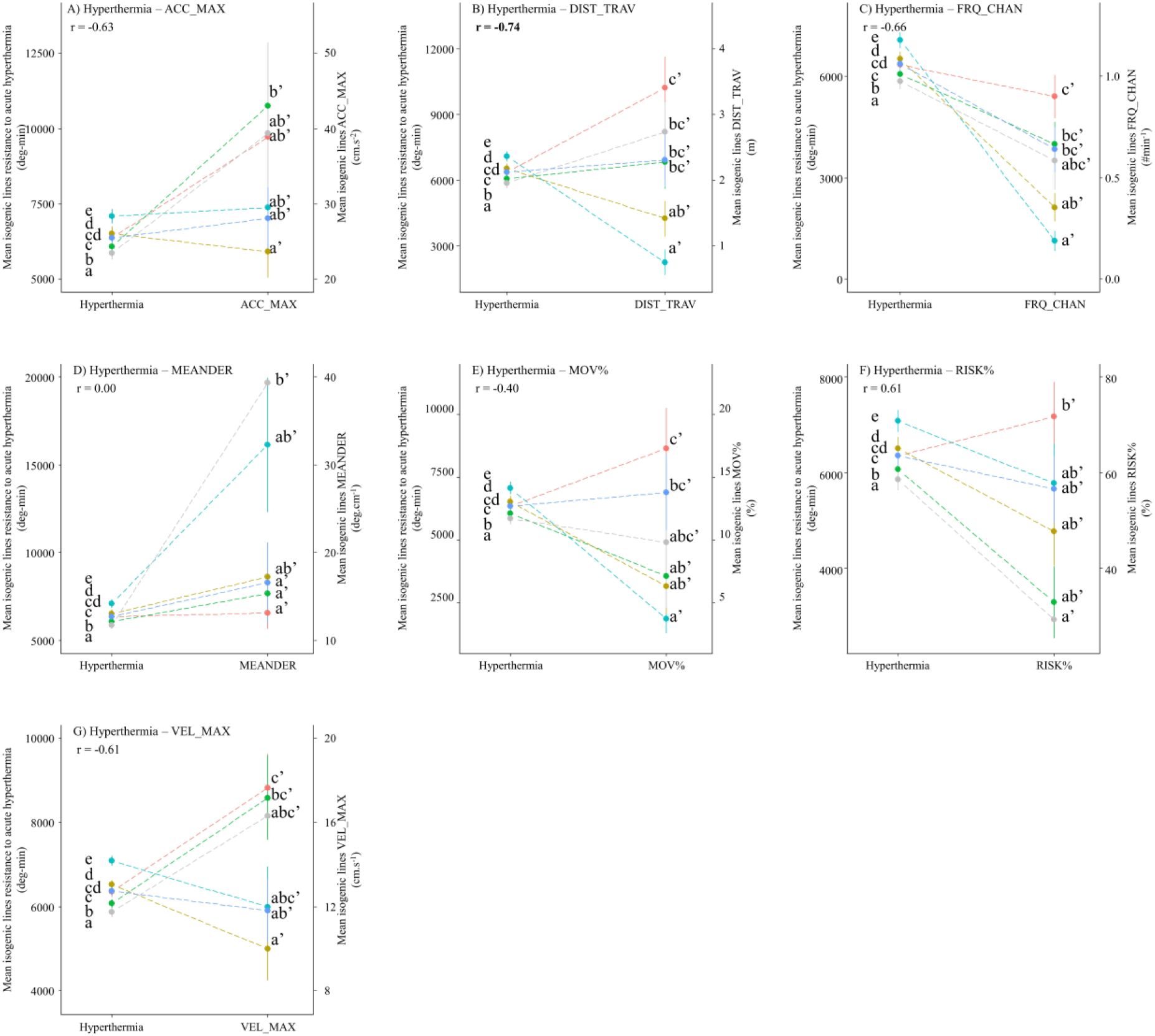
Relationships between mean isogenic line effects on behaviour traits and mean isogenic line effects on acute hyperthermia resistance. *r* represents the Pearson coefficient of correlation.

**Fig. 4.**
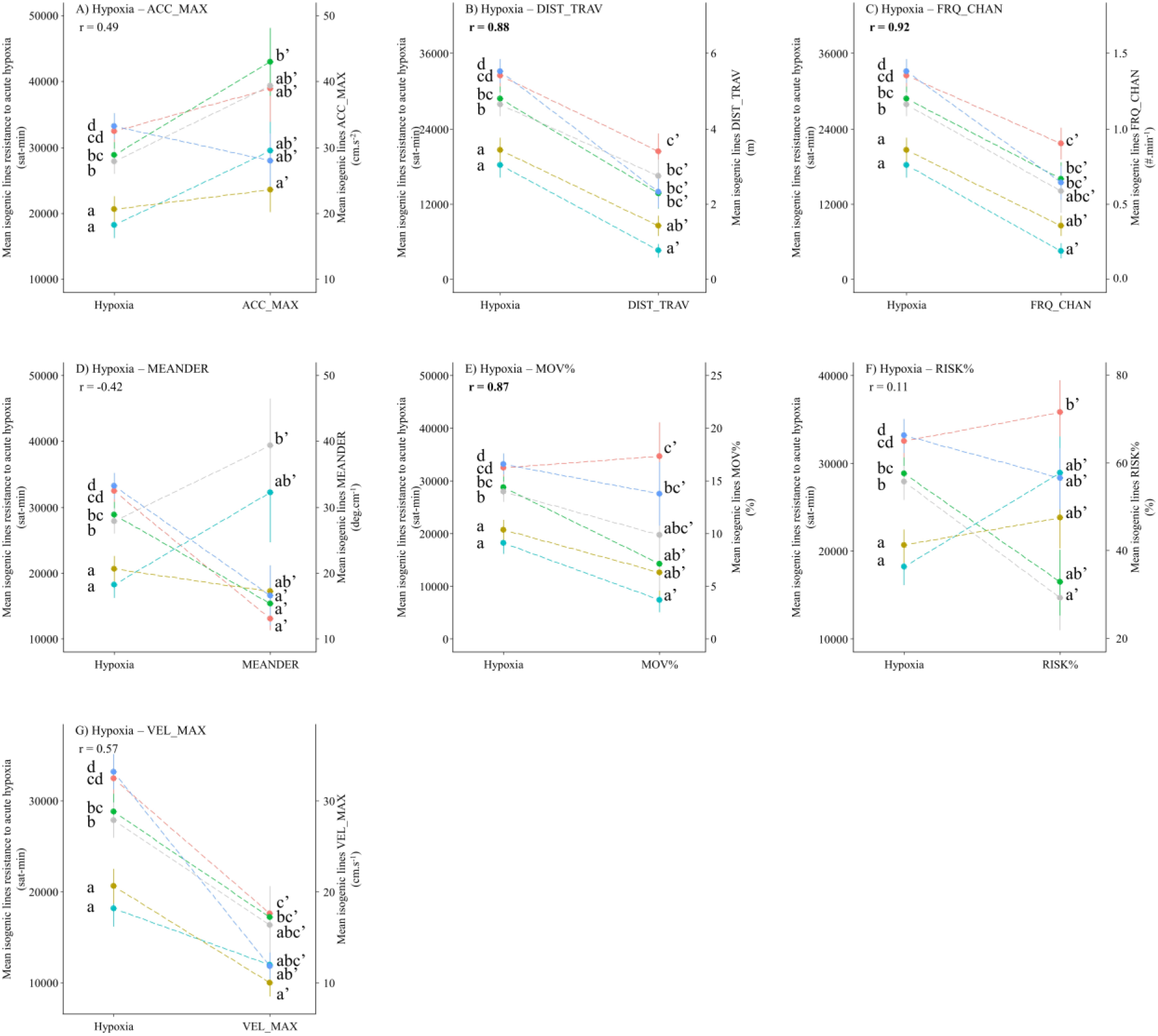
Relationships between mean isogenic line effects on behaviour traits and mean isogenic line effects on acute hypoxia resistance. *r* represents the Pearson coefficient of correlation.

However, despite these high correlations, it is important to note that the correlations between behaviour variables and resistance phenotypes were only significant for DIST_TRAV and hypoxia (p-value = 0.02), FRQ_CHAN and hypoxia (p-value = 0.01) and MOV% and hypoxia (p-value = 0.02). A part of the non-significant correlations is discussed hereafter as they possibly reveal relationships between behaviour variables and resistance phenotypes, which should be confirmed in future studies.

## 3 Discussion

In this work, we identified links between behaviour variables and resistance to acute hyperthermia e.g. travelling distance and frequency of change between the tanks’ zones or resistance to acute hypoxia e.g. travelling distance, frequency of change between the tanks’ zones and the percentage of time in movement.

### 3.1 Correcting resistance and behaviour phenotypes with body size

There were significant differences in body size between isogenic lines in the present experiments. Body size is a known component of the resistance to acute hyperthermia and hypoxia resistance (Perry et al., 2005; Nilsson & Östlund-Nilsson, 2008; McKenzie et al., 2021) and of the behaviour of fish (Millot et al., 2014). It was also the case in the present study as body size had a significant effect on resistance phenotypes and most behaviour variables. In the present paper, we decided to focus on the other components of the resistance to acute hyperthermia and hypoxia resistance and phenotypes were therefore corrected for body size effect.

### 3.2 Behaviours’ evolution with time and isogenic lines

The maximum acceleration and the maximum velocity significantly evolved during the behaviour test. At the beginning of the behaviour test, high maximum acceleration and velocity were observed, which is characteristic of bursts of high activity bouts behaviour. This enhanced activity could be linked to a high level of stress (Brown, Gardner & Braithwaite, 2005; Staven et al., 2019) and is consistent with the stress induced by the opening of the door and the novel environment proposed. It could also reveal the expression of a natural behaviour to quickly explore environment to escape stressful situations.

Significant behavioural differences between isogenic lines were found. These differences are consistent with the genetic component in fish behaviour, which was reviewed in Bell (2008), as well as reported in an earlier study that also used some rainbow trout isogenic lines (Millot et al., 2014). In particular, Millot et al. (2014) studied four isogenic lines that are investigated here: A02h, A22h, B45h and N38h. Establishing direct links between their results and ours is not simple since protocols and devices were different; specifically, Millot et al. used novel object and emergence tests. It is interesting to note that albeit evaluated in different contexts, A02h line travelled longer distances than B45h in both studies, carried out a decade ago, supporting the hypothesis of a genetic basis for these specific traits.

Behaviour variables reflecting the level of activity of fish (*i.e.* maximum acceleration, distance travelled, zone change frequency, maximum velocity) were highly correlated together at the isogenic line level. A strong correlation between activity variables was already found in rainbow trout for similar variables (Makaras et al., 2020).

### 3.3 Behaviour associated with acute hyperthermia or hypoxia resistance

Behavioural changes associated to challenging situation, e.g. hypoxia, are very variable between fish species and context (Chapman & Mckenzie, 2009) making challenging comparisons between studies.

Acute hyperthermia resistance appeared to be slightly associated with a low travelled distance and a low frequency of change between zones during the behaviour test. These results are consistent with the study of Campos, Val & Almeida-Val (2018), although the two studies are very different. They reported that fish species with the lowest routine activity level, indirectly measured with a proxy, had the highest resistance to acute hyperthermia. They also hypothesised that fish with the lowest activity have lower resting energetic demands and thus can resist longer acute hyperthermia (Campos, Val & Almeida-Val, 2018).

Acute hypoxia resistance was found to be strongly positively associated with the level of activity of fish in behaviour test. These results are difficult to compare to the literature. In rainbow trout exposed to sub-lethal levels of acute hypoxia, divergent behaviours were identified with some fish staying quiet and not moving while other fish actively tried to avoid hypoxia and showed peak activity (Van Raaij et al., 1996b). The survival rate of fish during the post-exposition recovery period was significantly associated with behavioural strategy, as fish staying quiet during the hypoxia challenge had a higher survival rate after return to normoxia compared to fish displaying high level of activity during the hypoxia challenge (Van Raaij et al., 1996b). In contrast, we showed in this study that isogenic lines resistant to acute hypoxia were rather highly active. The Van Raaij et al. experiment, performed with individuals from heterogeneous origins, established a link between behaviour during the hypoxia challenge and the later survival rate which is completely different from our experiments in which we established a link between hypoxia resistance and responses in a behavioural challenge performed in normoxia.

It is difficult to explain the underlying reasons of the observed association between acute hyperthermia and hypoxia resistances and behaviour variables because of an absence of consensus within the scientific community about the physiological mechanisms of the loss of equilibrium during acute hyperthermia or hypoxia stress (Nilsson & Östlund-Nilsson, 2008; Ern, Andreassen & Jutfelt, 2023). For acute hyperthermia resistance, the main theories about these mechanisms are the failure of short-term essential organs such as the brain, the failure of mitochondria processes, the thermal inactivation of enzymes, the desynchronization of interdependent metabolic processes because of different responses to temperature, the alteration of membrane structures, and a lack of oxygen (Fry, 1971; Schmidt-Nielsen, 1997; Pörtner, 2010; Chung & Schulte, 2020; Lefevre, Wang & McKenzie, 2021; Ern, Andreassen & Jutfelt, 2023). For hypoxia, loss of equilibrium during acute hypoxia challenge may be caused by an inability to produce adenosine triphosphate (ATP) fast enough with anaerobic substrate to compensate the failure of the aerobic metabolism, or by damage induced by the harmful metabolites produced by anaerobic metabolism, such as lactate or H+ (Nilsson & Östlund-Nilsson, 2008; Mandic, Todgham & Richards, 2009; Bergstedt, Pfalzgraff & Skov, 2021). However, it is still possible to make hypothesis about the relationship between the behaviour and the resistance phenotypes observed in the present paper. The weak pattern between low level of activity and resistance to acute hyperthermia could be interpreted as fish resisting better to acute hyperthermia are the ones saving energy by staying calm during the thermal challenge. However, the individual behaviour test was realised in normal temperature conditions and this relationship should thus be investigated at high temperature which is a well-known factor affecting fish behaviours such as schooling, activity, and maximum swimming speed (Hurst, 2007; Colchen et al., 2017; Kochhann et al., 2021; Pilakouta et al., 2022). A hypothesis about the strong pattern observed between high level of activity and resistance to acute hypoxia is that the most resistant fish to hypoxia are the ones that actively searched the areas in the challenge tank with the highest oxygen concentration, explaining their higher resistance. This kind of behaviour was observed in European sea bass, with individuals with higher metabolic rates showing greater hypoxia-induced increases in activity allowing them to find areas with higher oxygen concentration (Killen et al., 2012). Similarly, European catfish was shown in natural environment to actively search for the area with the highest O_2_ concentration in case of hypoxic conditions instead of limiting its movements to save energy and reduce O_2_ needs (Westrelin, Boulêtreau & Santoul, 2022). However, this hypothesis could only be valid in the case of the acute hypoxia challenge we conducted, if there was a O_2_ stratification in the tank during the challenge. We tried to avoid O_2_ stratification by maintaining a flow of water inside the tank during the challenge, but we did not check if it was effective. We recommend to measure O2 stratification in challenge tank for future hypoxic challenge. Assuming this hypothesis is valid, acute hypoxia resistance as it was phenotyped might not be adapted to improve fish resistance to hypoxic conditions in a farming environment. Indeed, in farming conditions there is no or little area with higher O_2_ concentration compared to natural conditions. Fish increasing their activity to find the areas with the higher O_2_ concentration might therefore be maladapted as this kind of behaviour decreases the O_2_ reserve of the tank faster and, therefore, reduces the survival probability of the whole batch of farmed fish. Further research on this subject would be important to check the relevance of acute hypoxia resistance in farming conditions.

### 3.4 Potential of behaviour as a proxy of acute hyperthermia and hypoxia resistance traits

In the present study, some behaviour variables were strongly associated with hypoxia resistance and moderately with acute hyperthermia resistance, at the level of the isogenic lines. These results suggest a potential for using behavioural variables as novel, non-stressful proxies for acute hyperthermia and hypoxia resistance in fish, improving experimental fish welfare.

However, as associations revealed with only six genotypes can give misleading results, these results still need to be validated on more genotypes and different populations. In case of confirmation, it would also be necessary to optimise how behaviour is measured, as data acquisition and analysis of the measurements are time-consuming compared to the classical phenotyping method. For instance, in (Lagarde et al., 2023b), it took around 700 minutes to phenotype the resistance to acute hyperthermia or hypoxia stress of 300 fish, equivalent to 0.4 fish phenotyped/minute, while in the present study, it took 25 minutes to phenotype six fish, equivalent to a rate of 0.2 phenotyped fish/minute. In addition, while the classical resistance phenotyping method directly gives the phenotypes of resistance, the behaviour phenotyping method implies extracting and analysing behaviour variables from recorded videos. However, the purpose of the behavioural phenotyping protocols in the present study was to identify candidate discriminating behaviours before starting optimisation. Future studies could implement significant time savings such as determining the minimum behaviour phenotyping time period required, increasing the number of fish in behaviour test, automating data extraction, etc. The accuracy of behaviour variables in discriminating phenotypes of resistance to acute hyperthermia and hypoxia must also be determined at the fish level to determine their value as a proxy.

### 3.5 Implications for selective breeding of acute hyperthermia and hypoxia resistance

The correlations found between some behaviour variables and acute hypoxia resistance, and potentially acute hyperthermia resistance, at the level of the isogenic line might reveal genetic correlations between some behavioural variables and resistance phenotypes. Therefore, selection for resistance to acute hyperthermia or hypoxia stress, as proposed by (Perry et al., 2005), (Lagarde et al., 2023a), (Prchal et al., 2023), could result in the indirect selection of certain behavioural variables. This side effect is not surprising since, in practice, selected fish over several generations are known to exhibit different behaviours, such as swimming, foraging, predator avoidance, social behaviour, personality, reproduction and cognitive learning, compared to the wild type (Pasquet, 2019). Specifically, for rainbow trout, domestication was shown to reduce predator avoidance behaviours (Johnsson, 1993; Berejikian, 1995). Behaviour modifications of farmed fish through direct or indirect genetic selection can positively or negatively impact fish adaptation to farming systems (Doyle & Talbot, 1986). If resistance to acute hyperthermia and hypoxia were to be selected in a breeding program, special consideration must be paid to the effects of selecting these resistance phenotypes on the behaviour variables. More generally, physiological mechanisms underlying acute hyperthermia and hypoxia resistance are still not known with certainty (Nilsson & Östlund-Nilsson, 2008; Ern, Andreassen & Jutfelt, 2023), and unintended trade-offs with important breeding traits could therefore arise.

## Conclusion

In the present study, we identified several behaviour variables strongly associated with acute hypoxia resistance and moderately with acute hyperthermia resistance in rainbow trout, at the line level. These results support the idea that behaviour phenotyping could be a non-invasive method for phenotyping acute hyperthermia or hypoxia resistances. These conclusions, notably the correlations between behaviour and resistance traits, must be verified in a higher number of isogenic lines and other populations of rainbow trout, as well as at the fish level.

## Ethical statement

Experiments were carried out at the INRAE experimental facilities (PEIMA, INRAE, 2021, Fish Farming systems Experimental Facility, doi: 10.15454/1.5572329612068406E12, Sizun, France) authorised for animal experimentation under the French regulation C29-277-02. Experiments were carried out according to the European guidelines; the protocols were evaluated and approved by the ethical committee CEFEA No 74 and authorised by the French Ministry of Higher Education and Research (APAFIS#24784_2020032510486957).

## Acknowledgements

This study was supported by the European Maritime and Fisheries Fund and FranceAgrimer (Hypotemp project, n° P FEA470019FA1000016). We are grateful to the Academic Writing Center of Université Paris Saclay for their valuable help in proofreading.

