## Supplementary Figures for "Screening for links between behaviour and acute hyperthermia and hypoxia resistance in rainbow trout using isogenic lines"

**Supplementary Fig. S1** Video capture of behaviour experimental device.

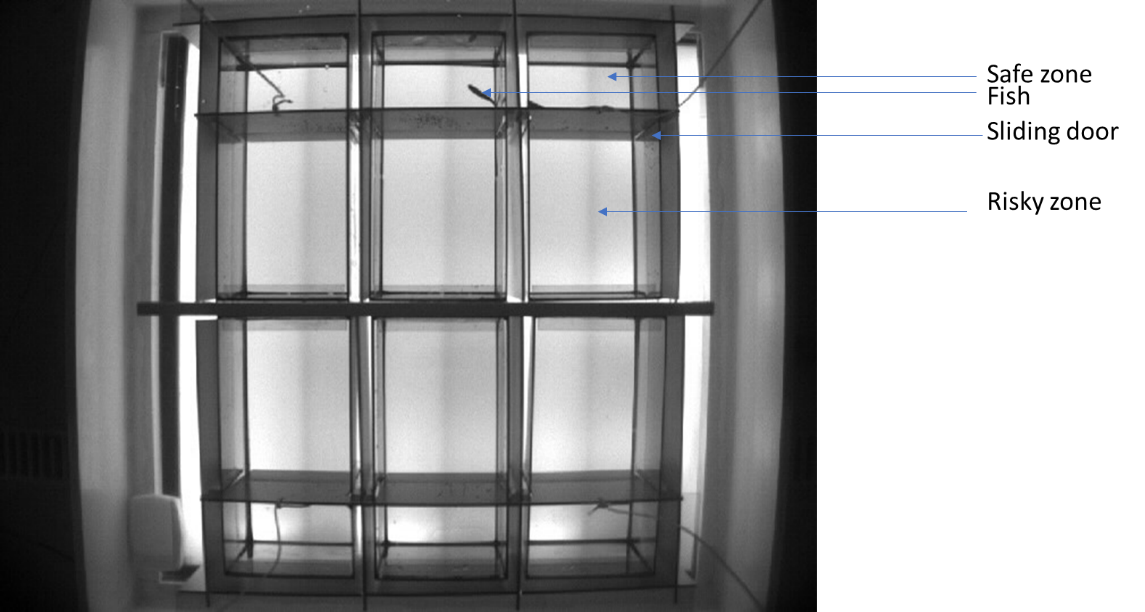

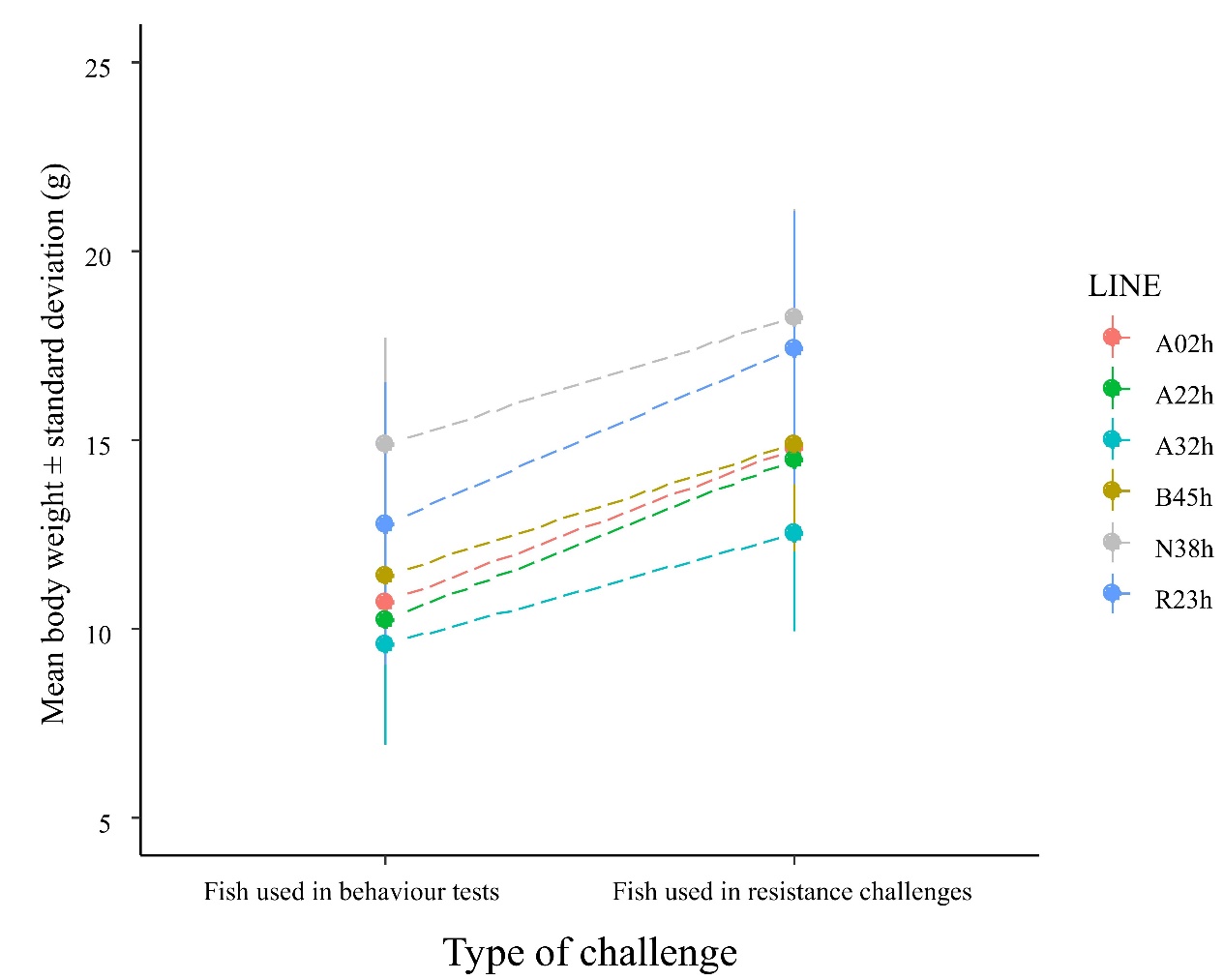
**Supplementary Fig. S2** Isogenic lines mean body weight of fish in behaviour tests and resistance challenges
