## Supplementary Tables for "Screening for links between behaviour and acute hyperthermia and hypoxia resistance in rainbow trout using isogenic lines"

**Supplementary Table S1** Mean body weight (g) of isogenic lines in behaviour test and resistance challenges. Means with same letters within columns are not significantly different. sd: standard deviation.

| Line | Behaviour test  mean body weight (± sd) | Resistance challenge  mean body weight (± sd) |
| --- | --- | --- |
| A02h | 10.7^a^ ± 2.0 | 14.8^b^ ± 2.8 |
| A22h | 10.3^a^ ± 2.8 | 14.5^b^ ± 2.7 |
| A32h | 9.6^a^ ± 2.6 | 12.5^a^ ± 2.6 |
| B45h | 11.4^ab^ ± 2.1 | 14.9^b^ ± 2.8 |
| N38h | 14.9^c^ ± 2.8 | 18.3^c^ ± 2.8 |
| R23h | 12.8^b^ ± 3.8 | 17.4^c^ ± 3.6 |

**Supplementary Table S2** Details about the significance of the fixed structure of the best model. In F-test, “*” means the *P*-value is inferior to 0.05 significant threshold.

| **Behaviour variables** | **Line** | **Time** | **Length** | **line*length** |
| --- | --- | --- | --- | --- |
| ACC_MAX | **F(5, 507) = 6.8*** | **F(4, 507) = 5.5*** | **F(1, 507) = 4.7*** | **F(5, 507) = 2.7*** |
| DIST_TRAV | **F(5, 516) = 8.4*** | NA | F(1, 516) = 0.1 | NA |
| EMERGENCE | NA | NA | NA | NA |
| FRQ_CHAN | **F(5, 516) = 10.5*** | NA | F(1, 516) = 0.3 | NA |
| MEANDER | **F(5, 289) = 3.7*** | NA | NA | NA |
| MOV% | **F(5, 511) = 6.5*** | NA | F(1, 511) = 1.7 | **F(5, 511) = 3.1*** |
| RISK% | **F(5, 511) = 9.0*** | NA | **F(1, 511) = 4.2*** | **F(5, 511) = 3.7*** |
| VEL_MAX | **F(5, 507) = 7.9*** | **F(4, 507) = 5.1*** | F(1,507) = 0.0 | **F(5, 507) = 2.3*** |

**Supplementary Table S3** Regression coefficients of *fork length* (cm) on behaviour variables. *fork length* regression coefficients for behaviour variables with significant interaction between *fork length* and *isogenic line* effect are given for each isogenic line.

| Isogenic lines | ACC_MAX (cm.s^-2^) | MOV% (%) | VEL_MAX (cm.s^-1^) | RISK% (%) |
| --- | --- | --- | --- | --- |
| A02h | -12 | -4,7 | -2,3 | 26,4 |
| A22h | -7,6 | -1,1 | -2,9 | -4,3 |
| A32h | 15,2 | 0,6 | 5,7 | 11 |
| B45h | -0,7 | -1,3 | -0,2 | 12,2 |
| N38h | 3 | -0,6 | -1 | -1,5 |
| R23h | 0,9 | -8,2 | -1,4 | -9,9 |

**Supplementary Table S4** Effect of *time bin* on behaviour variables for which *time bin* effect was significant. Means with same letters within columns are not significantly different. Time bins: 1 (0-5 minutes), 2 (5-10 minutes), 3 (10-15 minutes), 4 (15-20 minutes) and 5 (20-25 minutes).

| Time bin | ACC_MAX | VEL_MAX |
| --- | --- | --- |
|  | (cm.s-2) | (cm.s-1) |
| 1 | 46,3^b^ | 18,5^b^ |
| 2 | 33,7^ab^ | 14,7^ab^ |
| 3 | 25,7^a^ | 11,9^a^ |
| 4 | 35,9^ab^ | 13,6^a^ |
| 5 | 27,0^a^ | 12,1^a^ |
